# Genome-wide bioinformatic analyses predict key host and viral factors in SARS-CoV-2 pathogenesis

**DOI:** 10.1101/2020.07.28.225581

**Authors:** Mariana G. Ferrarini, Avantika Lal, Rita Rebollo, Andreas Gruber, Andrea Guarracino, Itziar Martinez Gonzalez, Taylor Floyd, Daniel Siqueira de Oliveira, Justin Shanklin, Ethan Beausoleil, Taneli Pusa, Brett E. Pickett, Vanessa Aguiar-Pulido

**Affiliations:** University of Lyon, INSA-Lyon, INRA, BF2I, Villeurbanne, France; NVIDIA Corporation, Santa Clara, CA, USA; Oxford Big Data Institute, Nuffield Department of Medicine, University of Oxford, Oxford, UK; Centre for Molecular Bioinformatics, Department of Biology, University Of Rome Tor Vergata, Rome, Italy; Amsterdam UMC, Amsterdam, The Netherlands; Center for Neurogenetics, Weill Cornell Medicine, Cornell University, New York, NY, USA; Laboratoire de Biométrie et Biologie Evolutive, Université de Lyon; Université Lyon 1; CNRS; UMR 5558, Villeurbanne, France; Brigham Young University, Provo, UT, USA; Luxembourg Centre for Systems Biomedicine (LCSB), University of Luxembourg, Belvaux, Luxembourg

**Keywords:** SARS-CoV-2, COVID-19, gene expression, RNA-seq, RNA-binding proteins, host-pathogen interaction, transcriptomics

## Abstract

The novel betacoronavirus named Severe Acute Respiratory Syndrome Coronavirus 2 (SARS-CoV-2) caused a worldwide pandemic (COVID-19) after initially emerging in Wuhan, China. Here we applied a novel, comprehensive bioinformatic strategy to public RNA sequencing and viral genome sequencing data, to better understand how SARS-CoV-2 interacts with human cells. To our knowledge, this is the first meta-analysis to predict host factors that play a specific role in SARS-CoV-2 pathogenesis, distinct from other respiratory viruses. We identified differentially expressed genes, isoforms and transposable element families specifically altered in SARS-CoV-2 infected cells. Well-known immunoregulators including *CSF2, IL-32, IL-6* and *SERPINA3* were differentially expressed, while immunoregulatory transposable element families were overexpressed. We predicted conserved interactions between the SARS-CoV-2 genome and human RNA-binding proteins such as hnRNPA1, PABPC1 and eIF4b, which may play important roles in the viral life cycle. We also detected four viral sequence variants in the spike, polymerase, and nonstructural proteins that correlate with severity of COVID-19. The host factors we identified likely represent important mechanisms in the disease profile of this pathogen, and could be targeted by prophylactics and/or therapeutics against SARS-CoV-2.

**Graphical Abstract:** 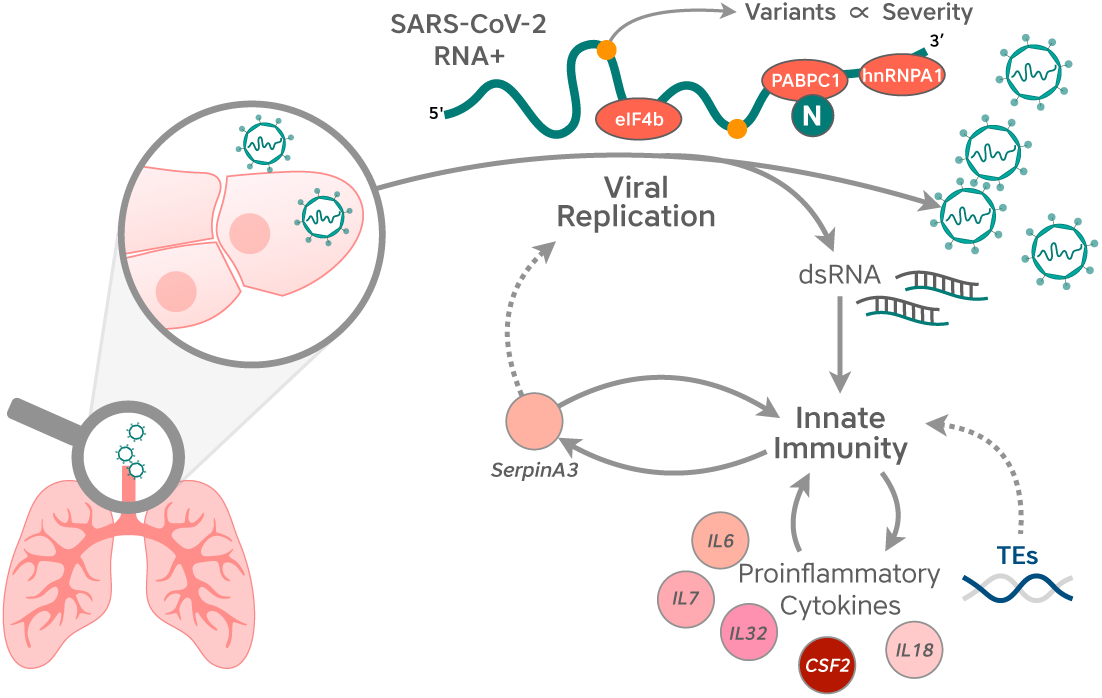

## Introduction

In December of 2019 a novel betacoronavirus that was named Severe Acute Respiratory Syndrome Coronavirus 2 (SARS-CoV-2) emerged in Wuhan, China [4, 29]. This virus is responsible for causing the coronavirus disease of 2019 (COVID-19) and, by August 18 of 2020, it had already infected more than 21 million people worldwide, accounting for at least 750 thousand deaths (https://covid19.who.int). The SARS-CoV-2 genome is phylogenetically distinct from the SARS-CoV and Middle East Respiratory Syndrome (MERS) betacoronaviruses that caused human outbreaks in 2002 and 2012 respectively [79, 87]. Based on its high sequence similarity to a coronavirus isolated from bats [88], SARS-CoV-2 is hypothesized to have originated from bat coronaviruses, potentially using pangolins as an intermediate host before infecting humans [39].

SARS-CoV-2 infects human cells by binding to the angiotensin-converting enzyme 2 (ACE2) receptor [85]. Recent studies have sought to understand the molecular interactions between SARS-CoV-2 and infected cells [24], unfortunately, only a few have quantified gene expression changes in patient samples or cultured lung-derived cells infected by this virus [10, 46, 83]. These studies are essential to understanding the mechanisms of pathogenesis and immune response which can facilitate the development of treatments for COVID-19 [35, 54, 89].

Viruses generally trigger a drastic host response during infection. A subset of these specific changes in gene regulation is associated with viral replication, and therefore can pinpoint potential drug targets. In addition, transposable element (TE) overexpression has been observed upon viral infection [50], and TEs have been actively implicated in gene regulatory networks related to immunity [15]. Moreover, SARS-CoV-2 is a virus with a positive-sense, single-stranded, monopartite RNA genome. Such viruses are known to co-opt host RNA-binding proteins (RBPs) for diverse processes including viral replication, translation, viral RNA stability, assembly of viral protein complexes, and regulation of viral protein activity [22, 45].

In this work we identified a signature of altered gene expression that is consistent across published datasets of SARS-CoV-2 infected human lung cells. We present extensive results from functional analyses (signaling pathway enrichment, biological functions, transcript isoform usage, metabolic flux prediction, and TE overexpression) performed upon the genes that are differentially expressed during SARS-CoV-2 infection [10]. We also predict specific interactions between the SARS-CoV-2 RNA genome and human RBPs that may be involved in viral replication, transcription or translation, and identify viral sequence variations that are significantly associated with increased pathogenesis in humans. Knowledge of these molecular and genetic mechanisms is important to understand the SARS-CoV-2 pathogenesis and to improve the future development of effective prophylactic and therapeutic treatments.

## Results

We designed a comprehensive bioinformatics workflow to identify relevant host-pathogen interactions using a complementary set of computational analyses (Figure 1). First, we carried out an exhaustive analysis of differential gene expression in the only publicly available dataset to date from human lung cells infected with SARS-CoV-2 along with other respiratory viruses. We identified gene, isoform- and pathway-level responses that specifically characterize SARS-CoV-2 infection. Second, we predicted putative interactions between the SARS-CoV-2 RNA genome and human RBPs. Th we identified a subset of these human RBPs which were also differentially expressed in response SARS-CoV-2. Finally, we predicted four viral sequence variants that could play a role in diseas severity.

**Figure 1.**
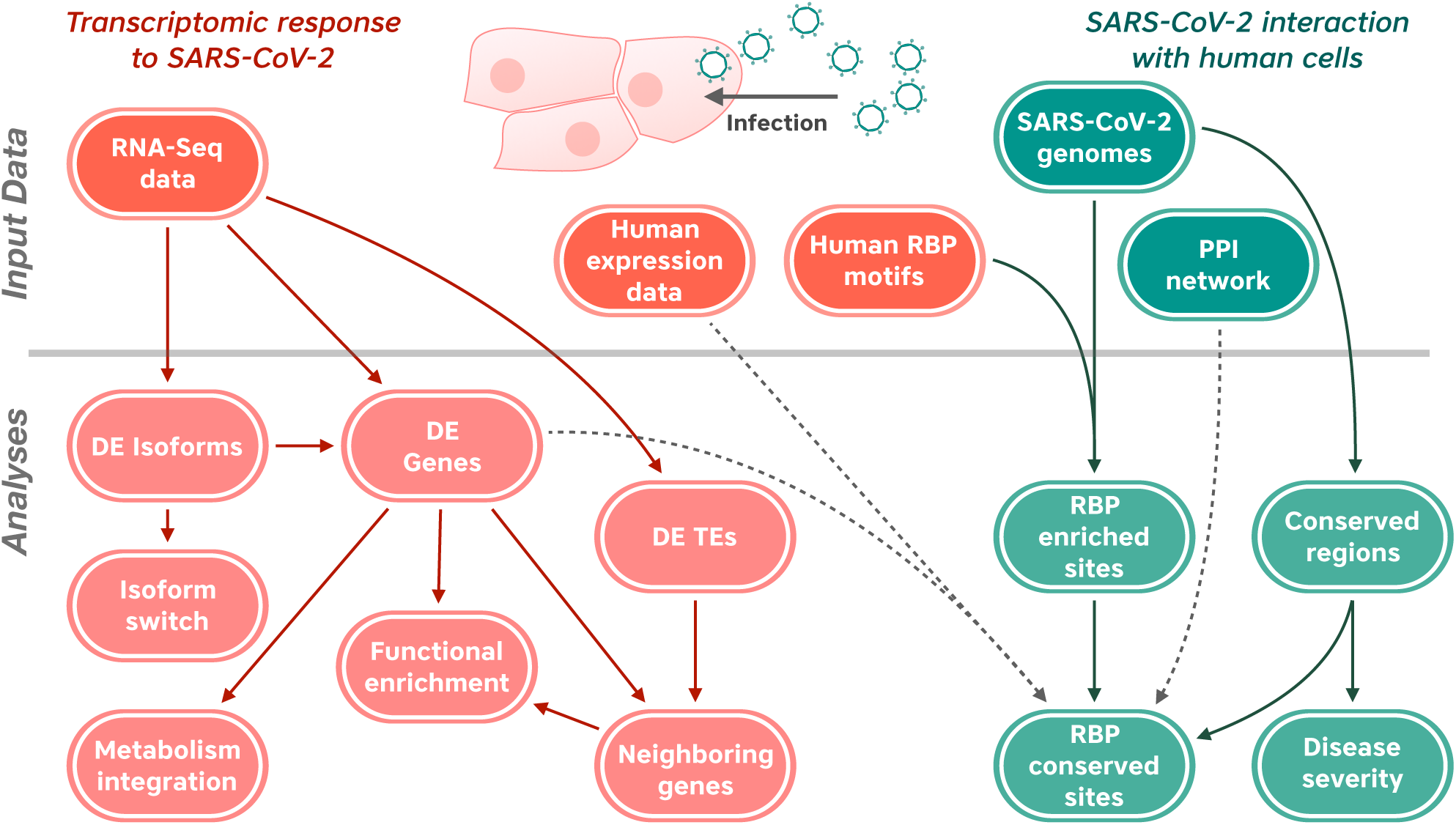
Overview of the bioinformatic workflow applied in this study.

### SARS-CoV-2 infection elicits a specific gene expression and pathway signature in human cells

We wanted to identify genes that were differentially expressed across multiple SARS-CoV-2 infected samples and not in samples infected with other respiratory viruses. As a primary dataset, we selected GSE147507 [10], which includes gene expression measurements from three cell lines derived from the human respiratory system (NHBE, A549, Calu-3) infected either with SARS-CoV-2, influenza A virus (IAV), respiratory syncytial virus (RSV), or human parainfluenza virus 3 (HPIV3), with different multiplicity of infection (MOI). We also analyzed an additional dataset GSE150316, which includes RNA-seq extracted from formalin fixed, paraffin embedded (FFPE) histological sections of lung biopsies from COVID-19 deceased patients and healthy individuals (see Figure 2A and Materials and Methods for further details).

**Figure 2.**
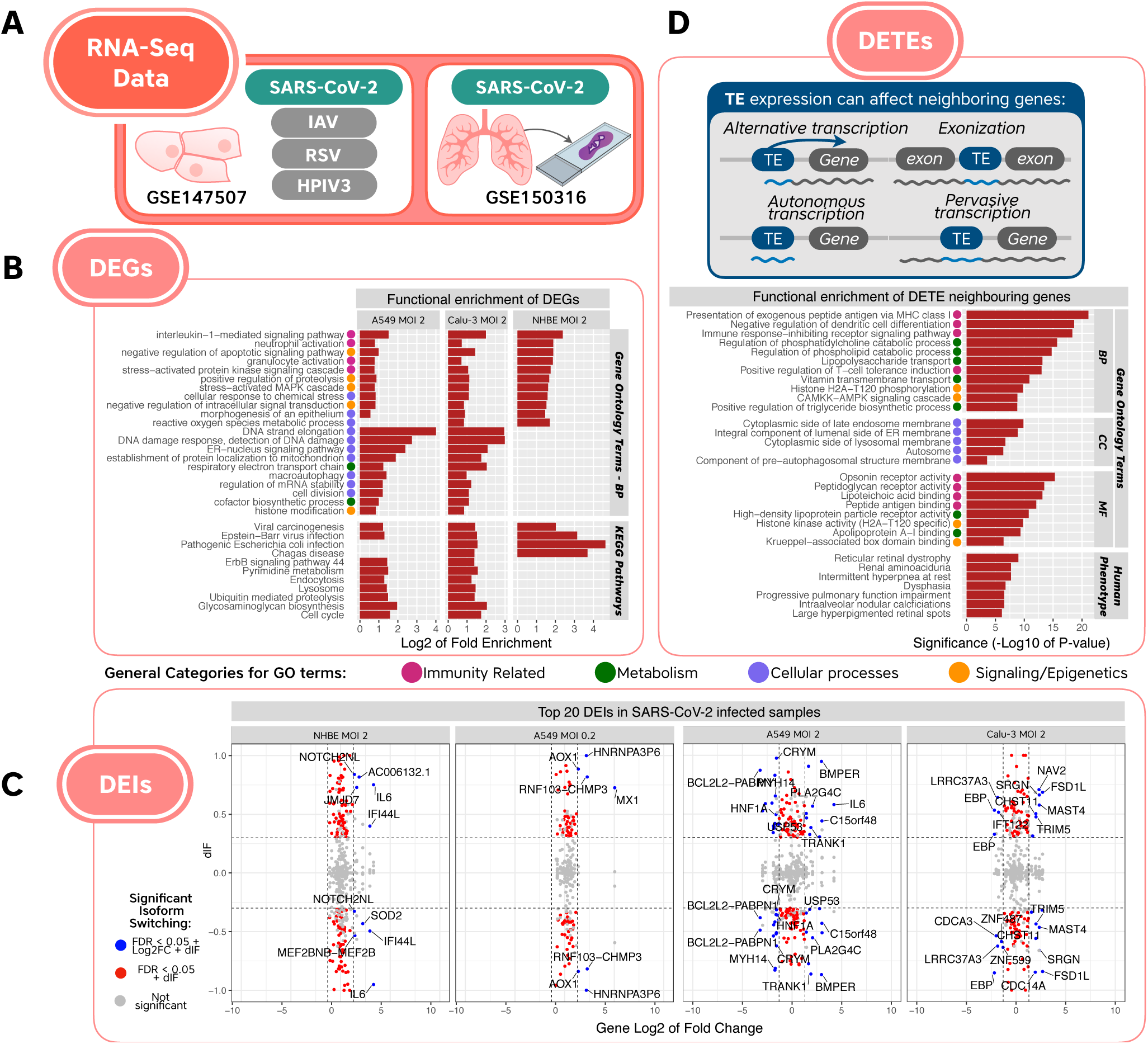
Overview of the RNA-seq based results specific to SARS-CoV-2 which were not detected in the other viral infections (IAV, HPIV3 and RSV). (A) Representation of the RNA-seq studies used in our analyses. (B) Non-redundant functional enrichment of DEGs. Here we report a subset of non-redundant reduced terms consistently enriched in more than one SARS-COV-2 cell line which were not detected in the other viruses’ datasets. We added generic categories of immunity, metabolism, cellular processes and signaling/epigeneitcs to the GO terms as colored dots. (C) Top 20 differentially expressed isoforms (DEIs) in SARS-CoV-2 infected samples. Y-axis denotes the differential usage of isoforms (dIF) whereas x-axis represents the overall log2FC of the corresponding gene. Thus, DEIs also detected as DEGs by this analysis are depicted in blue. (D) The upper right diagram depicts different manners by which TE family overexpression might be detected. While TEs may indeed be autonomously expressed, the old age of most TEs detected points toward either being part of a gene (exonization or alternative promoter), or a result of pervasive transcription. We report the functional enrichment for neighboring genes of differentially expressed TEs (DETEs) specifically upregulated in SARS-CoV-2 Calu-3 and A549 cells (MOI 2). The same categories used in subfigure (B) were attributed to the GO terms reported here.

Hence, we retrieved 41 differentially expressed genes (DEGs) that showed significant and consistent expression changes in at least three datasets from cell lines infected with SARS-CoV-2, and that were not significantly affected in cell lines infected with other viruses within the same dataset (Supplementary Table 1A). To these, we added 23 genes that showed significant and consistent expression changes in two of four cell line datasets infected with SARS-CoV-2 and at least one lung biopsy sample from a SARS-CoV-2 patient. Results coming from FFPE sections were less consistent presumably due to the collection of biospecimens from different sites within the lung. Thus, the final set consisted of 64 DEGs: 48 up-regulated and 16 downregulated of which 38 had an absolute Log2FC > 1 in at least one dataset (relevant genes from this list are shown in Table 1).

**Table 1.** Log2FC for selected genes that showed significant up-or down-regulation in SARS-CoV-2 infected samples (FDR-adjusted p-value < 0.05), and not in samples infected with the other viruses tested. Log2FC values are only provided for statistically significant samples.

| Gene | Cell Type and MOI |  |  |  | Biopsies |  |
| --- | --- | --- | --- | --- | --- | --- |
|  | A549<br>MOI 0.2 | A549<br>MOI 2 | Calu-3<br>MOI 2 | NHBE | Case<br>1 | Case<br>3 |
| *VNN2* |  | 6.18 | 0.42 | 6.13 |  |  |
| *CSF2* |  | 3.56 | 7.30 | 2.70 |  |  |
| *WNT7A* |  | 4.99 | 0.79 | 0.45 |  |  |
| *PDZK1IP1* |  | 1.72 | 0.70 | 2.28 |  |  |
| *SERPINA3* | 0.49 | 1.39 | 0.77 | 1.44 |  |  |
| *RHCG* |  | 1.51 | 2.02 | 1.33 | 2.53 |  |
| *IL32* |  | 1.64 | 1.23 | 1.21 |  |  |
| *PDGFB* |  | 1.91 | 1.75 | 1.00 |  |  |
| *ALDH1A3* |  | 1.09 | 1.32 | 0.39 |  |  |
| *TLR2* |  | 1.63 | 0.89 | 0.84 |  |  |
| *G0S2* |  | 0.66 | 3.79 | 0.83 |  |  |
| *NRCAM* |  | 0.73 | 1.82 | 0.78 |  |  |
| *SERPINB1* |  | 0.61 | 1.17 | 0.72 |  |  |
| *PRDM1* |  | 0.82 | 3.49 | 0.59 |  |  |
| *MT-TN* |  | 0.55 | 1.70 | 0.33 |  |  |
| *ATF4* |  | 0.79 | 1.07 | 0.26 |  |  |
| *BHLHE40* |  | 0.75 | 1.56 | 0.18 |  |  |
| *PTPN12* | 0.48 | 0.97 | 1.23 |  |  |  |
| *GPCPD1* | 0.36 | 0.94 | 1.69 |  |  |  |
| *DUSP16* | 0.33 | 0.41 | 1.43 |  |  |  |
| *FKBP5* |  | -0.39 |  | -0.36 | -1.47 | -2.14 |
| *DAP* |  | -0.18 | -0.61 |  | -1.16 |  |
| *FECH* |  | -0.27 | -0.36 |  | -1.54 |  |
| *MT-CYB* |  | -0.30 | -0.26 |  | -3.68 |  |
| *EIF4A1* |  | -0.33 | -0.63 |  | -1.85 |  |
| *POLE4* | -0.23 | -0.82 | -1.24 |  |  |  |
| *DDX39A* | -0.23 | -1.27 | -0.54 |  |  |  |
| *CENPP* |  | -0.36 | -0.40 | -0.38 |  |  |
| *TMEM50B* |  | -0.48 | -0.59 | -0.53 |  |  |
| *HPS1* | -0.28 | -0.31 | -0.62 |  |  |  |
| *SNX8* | -0.30 | -0.43 | -0.56 |  |  |  |

*SERPINA3*, an antichymotrypsin which was proposed as an interesting candidate for the inhibition of viral replication [13], was the only gene specifically upregulated in the four cell line datasets tested (Table 1). Other interesting up-regulated genes were the amidohydrolase *VNN2*, the pro-fibrotic gene *PDGFB*, the beta-interferon regulator *PRDM1* and the proinflammatory cytokines *CSF2* and *IL-32. FKBP5*, a known regulator of NF-kB activity, was among the consistently downregulated genes. We also generated additional lists of DEGs that met different filtering criteria (Supplementary Table 1B, see Supplementary File 1 for the complete DEG results for each dataset).

In order to better understand the underlying biological functions and molecular mechanisms associated with the observed DEGs, we performed a hypergeometric test to detect statistically significant overrepresented gene ontology (GO) terms among the DEGs having an absolute Log2FC > 1 in each dataset separately.

Consistent with the findings of Blanco-Melo et al. [10], GO enrichment analysis returned terms associated with immune system processes, Pi3K/AKT signaling pathway, response to cytokine, stress and virus, among others (see Supplementary File 2 for complete results). In addition, we report 285 GO terms common to at least two cell line datasets infected with SARS-CoV-2, and absent in the response to other viruses (Figure 2B, Supplementary Table 2A), including neutrophil and granulocyte activation, interleukin-1-mediated signaling pathway, proteolysis, and stress activated signaling cascades.

Next, we wanted to pinpoint intracellular signaling pathways that may be modulated specifically during SARS-CoV-2 infection. A robust signaling pathway impact analysis (SPIA) enabled us to identify 30 pathways, including many involved in the host immune response, that were significantly enriched among differentially expressed genes in at least one virus-infected cell line dataset (Supplementary Table 3). More importantly, we predicted four pathways to be specific to SARS-CoV-2 infection and observed that the significant pathways differed by cell type and multiplicity of infection. The significant results included only one term common to A549 (MOI 0.2) and Calu-3 cells (MOI 2), namely the interferon alpha/beta signaling. Additionally, we found the amoebiasis (A549 cells, MOI 0.2), the p75(NTR)-mediated and the trka receptor signaling pathways (A549 cells, MOI 2) as significantly impacted.

We also used a classic hypergeometric method as a complementary approach to our SPIA pathway enrichment analysis. While there were generally higher numbers of significant results using this method, we observed that the vast majority of enriched terms (FDR < 0.05) described infections with various pathogens, innate immunity, metabolism, and cell cycle regulation (Supplementary Table 3). Interestingly, we were able to detect enriched KEGG pathways common to at least two SARS-CoV-2 infected cell types and absent from the other virus-infected datasets (Figure 2B, Supplementary Table 2B). These included pathways related to infection, cell cycle, endocytosis, signalling pathways, cancer and other diseases.

### SARS-CoV-2 infection results in altered lipid-related metabolic fluxes

To integrate the gene expression changes with metabolic activity in response to virus infection, we projected the transcriptomic data onto the human metabolic network [77]. This analysis detected common decreased fluxes in inositol phosphate metabolism in both A549 and Calu-3 cells infected with SARS-CoV-2 at a MOI of 2 (Supplementary Table 4). The consensus solution (obtained taking into account the enumeration of all solutions) in A549 cells (MOI 2) also recovered decreased fluxes in several lipid pathways: fatty acid, cholesterol, sphingolipid, and glycerophospholipid. In addition, we detected an increased flux common to A549 and Calu-3 cell lines in reactive oxygen species (ROS) detoxification, in accordance with previous terms recovered from functional enrichment analyses.

### SARS-CoV-2 infection induced an isoform switch of genes associated with immunity and mRNA processing

We wanted to analyze changes in transcript isoform expression and usage associated with SARS-CoV-2 infection, as well as to predict whether these changes might result in altered protein function. We identified isoforms experiencing a switch in usage greater than or equal to 30% in absolute value, and retrieved those with a Bonferroni-adjusted p-value less than 0.05. After calculating the difference in isoform usage (dIF) per gene (in each condition), we performed predictive functional consequence and alternative splicing analyses for all isoforms globally as well as at the individual gene level.

We observed 3,569 differentially expressed isoforms (DEIs) across all samples (Supplementary Figure 1A, Supplementary Table 5A). Results indicate that isoforms from A549 cells infected with RSV, IAV and HPIV3 exhibited significant differences in biological events such as complete open reading frame (ORF) loss, shorter ORF length, intron retention gain and decreased sensitivity to nonsense mediated decay (Supplementary Figure 1B). These conditions also displayed various changes in splicing patterns, ranging from loss of exon skipping events, changes in usage of alternative transcription start and termination sites, and decreased alternative 5’ and 3’ splice sites (Supplementary Figure 1C).

In contrast, isoforms from SARS-CoV-2 infected samples displayed no significant global changes in biological consequences or alternative splicing events between conditions (Supplementary Figures 1A and 1B respectively). Trends indicated transcripts in SARS-CoV-2 samples experienced decreases in ORF length, numbers of domains, coding capability, intron retention and nonsense mediated decay (Supplementary Figure 1A). These biological consequences may result from increased multiple exon skipping events and alternative transcription start sites via alternative 5’ acceptor sites (Supplementary Figure 1B). While not significant, these trends implicate that the SARS-CoV-2 virus may globally trigger host cell machinery to generate shorter isoforms that, while not shuttled for degradation, either do not produce functional proteins or produce alternative aberrant proteins not utilized in non-SARS-CoV-2 tissue conditions.

Despite the lack of global biological consequence and splicing changes, individual isoforms from SARS-CoV-2 infected samples experienced significant changes in gene expression and isoform usage (Figure 2C). Top-expressing genes were associated with cellular processes such as immune response and antiviral activity (*IFI44L, IL6, MX1, TRIM5*), transcription and mRNA processing (*DDX10, HNRNPA3F6, JMJD7, ZNF487, ZNF599*) and cell cycle and survival (*BCL2L2-PABPN1, CDCA3*) (Supplementary Table 5B). Similarly, significant genes from non-SARS-CoV-2 samples were associated with processes such as immune cell development and response (*ADCY7, BATF2, C9orf72, ETS1, GBP2, IFIT3*), transcription regulation and DNA repair (*ABHD14B, ATF3, IFI16, POLR2J2, SMUG1, ZNF19, ZNF639*), mitochondrial function (*ATP5E, BCKDH8, TST, TXNRD2*), and GTPase activity (*GBP2, RAP1GAP, RGS20, RHOBTB2*) (Supplementary Figure 1D, Supplementary Table 5B).

Upon further inspection, we noticed that *IL-6*, a gene encoding a cytokine involved in acute and chronic inflammatory responses, displayed 3 and 4-fold increases in expression in NHBE and A549 cells, respectively (infected with a MOI of 2) (Supplementary Figure 1B). To date, the Ensembl Genome Reference Consortium has identified 9 *IL-6* isoforms in humans, with the traditional transcript having 6 exons (*IL6-204*), 5 of which contain coding elements. NHBE cells expressed 4 known *IL-6* isoforms, while A549 cells expressed 1 unknown and 6 known isoforms. When evaluating the actual isoforms used across conditions, NHBE cells used 3 out of 4 isoforms observed, while A549 cells used all 7 observed isoforms. Isoform usage is evaluated based on isoform fraction (IF), or the percentage of an isoform found relative to all other identified isoforms associated with a specific gene. For example, in the case of NHBE SARS-CoV-2 samples, the IF for the *IL6-201* isoform = 0.75, *IL6-204* = 0.05, *IL6-206* = 0.09, *IL6-209* = 0.06, and the sum of these IF values = 0.95, or 95% usage of the *IL-6* gene. Both SARS-CoV-2 samples exhibited exclusive usage of non-canonical isoform *IL6-201*, and inversely, mock samples almost exclusively utilized the *IL6-204* transcript. In NHBE infected cells, isoform *IL6-201* experienced a significant increase in usage (dIF = 0.75) and *IL6-204* a significant decrease in usage (dIF = -0.95) when compared to mock conditions. Similarly, isoform *IL6-201* in A549 infected cells experienced an increase in usage (dIF = 0.58), while uses of all other isoforms remained non-significant in comparison to mock conditions.

### Overexpression of TE families close to immune-related genes upon SARS-CoV-2 infection

In order to estimate the expression of TE families and their possible roles in SARS-CoV-2 infection, we mapped the RNA-seq reads against all annotated human TE families and detected DETEs (Supplementary File 3). We found 68 common TE families upregulated in SARS-CoV-2 infected A549 and Calu-3 cells (MOI 2). From this list, we excluded all TE families detected in A549 cells infected with the other viruses. This allowed us to identify 16 families that were specifically upregulated in Calu-3 and A549 cells infected with SARS-CoV-2 and not in the other viral infections.

The 16 families identified were MER77B, MamRep4096, MLT2C2, PABL A, Charlie9, MER34A, L1MEg1, LTR13A, L1MB5, MER11C, MER41B, LTR79, THE1D-int, MLT1I, MLT1F1, MamRep137. Most of the TE families uncovered are ancient elements, incapable of transposing, or harboring ntrinsic regulatory sequences [37, 57, 70]. Eleven of the 16 TE families specifically upregulated in SARS-COV-2 infected cells are long terminal repeat (LTR) elements, and include well known TE immune regulators. For instance, the MER41B (primate specific TE family) is known to contribute to Interferon gamma inducible binding sites (bound by STAT1 and/or IRF1) [14, 66].

Other LTR elements are also enriched in STAT1 binding sites (MLT1L) [14], or have been shown to act as cellular gene enhancers (LTR13A [16, 32]).

Given the propensity for the TE families detected to impact nearby gene expression, we further investigated the functional enrichment of genes near upregulated TE families (+- 5kb upstream, 1kb downstream). We detected GO functional enrichment of several immunity-related terms (e.g. MHC protein complex, antigen processing, regulation of dendritic cell differentiation, T-cell tolerance induction), metabolism related terms (such as regulation of phospholipid catabolic process), and more interestingly a specific human phenotype term called “Progressive pulmonary function impairment” (Figure 2D). Even though we did not limit our search only to neighboring genes which were also DE, we found several similar (and very specific) enriched terms in both analyses, for instance related to immune response, endosomes, endoplasmic reticulum, vitamin (cofactor) metabolism, among others. This result supports the idea that some responses during infection could be related to TE-mediated transcriptional regulation. Finally, when we searched for enriched terms related to each one of the 16 families separately, we also detected immunity related enriched terms such as regulation of interleukins, antigen processing, TGFB receptor binding and temperature homeostasis (Supplementary File 4). It is important to note that given the old age of some of the TEs detected, overexpression might be associated with pervasive transcription, or inclusion of TE copies within unspliced introns (see upper box in Figure 2D).

### The SARS-CoV-2 genome is enriched in binding motifs for 40 human RBPs, most of them conserved across SARS-CoV-2 genome isolates

Our next aim was to predict whether any host RNA binding proteins interact with the viral genome. To do so, we first filtered the AtTRACT database [23] to obtain a list of 102 human RBPs and 205 associated Position Weight Matrices (PWMs) describing the sequence binding preferences of these proteins. We then scanned the SARS-CoV-2 reference genome sequence to identify potential binding sites for these proteins. Figure 3 illustrates our analysis pipeline.

**Figure 3.**
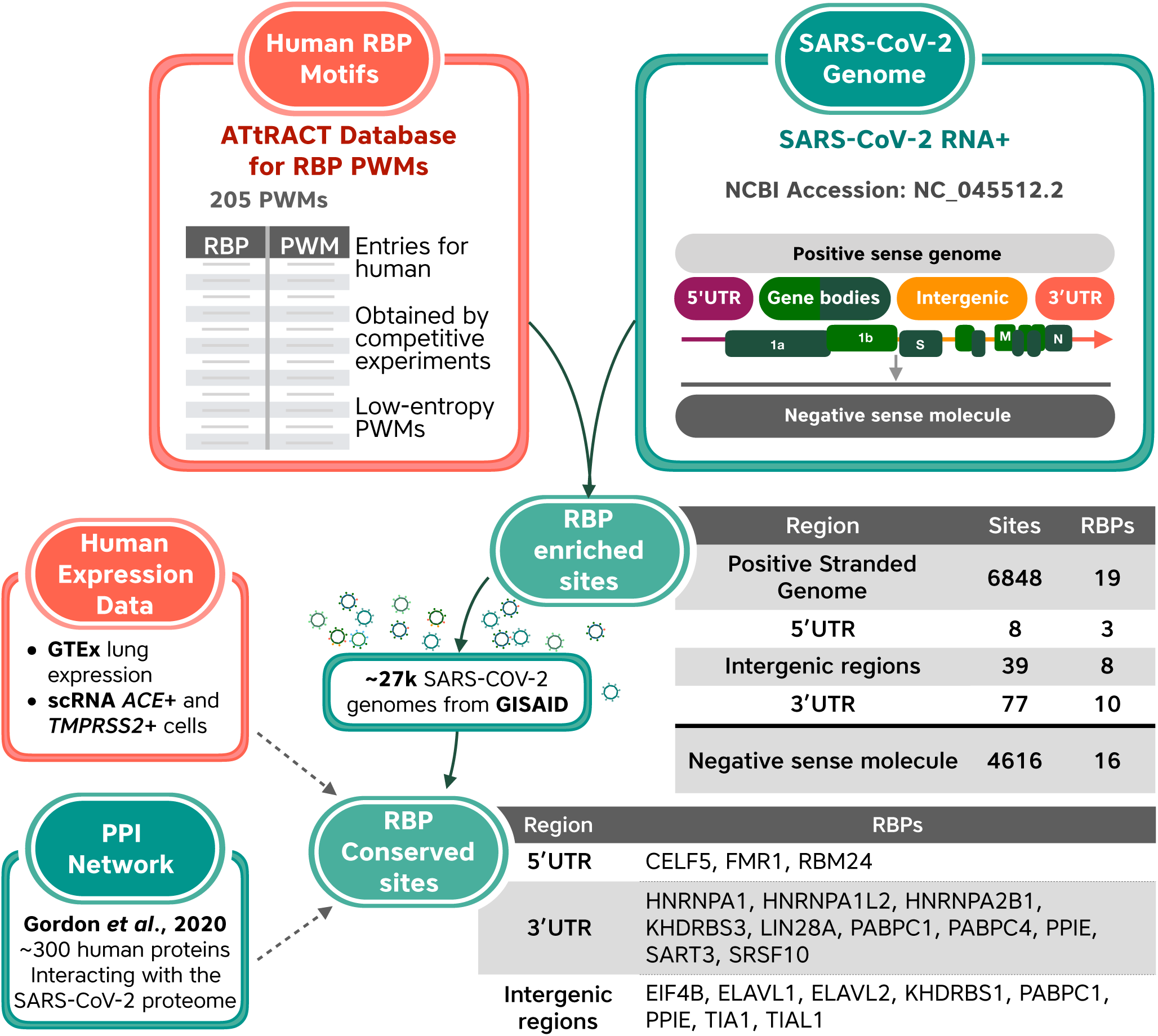
Workflow and selected results for analysis of potential binding sites for human RNA-binding proteins in the SARS-CoV-2 genome.

We identified 99 human RBPs with 11,897 potential binding sites in the SARS-CoV-2 positive-sense genome. Since the SARS-CoV-2 genome produces negative-sense intermediates as part of the replication process [36], we also scanned the negative-sense molecule, where we found 11,333 potential binding sites for 96 RBPs (Supplementary Table 6).

To find RBPs whose binding sites occur in the SARS-CoV-2 genome more often than expected by chance, we repeatedly scrambled the genome sequence to create 1,000 simulated genome sequences with an identical nucleotide composition to the SARS-CoV-2 genome sequence (30% A, 18% C, 20% G, 32% T). We used these 1,000 simulated genomes to determine a background distribution of the number of binding sites found for a specific RBP. This allowed us to pinpoint RBPs with significantly more or significantly fewer binding sites in the actual SARS-CoV-2 genome than expected based on the background distribution (two-tailed z-test, FDR-corrected P < 0.01). To retrieve RBPs whose motifs were enriched in specific genomic regions, we also repeated this analysis independently for the SARS-CoV-2 5’UTR, 3’UTR, intergenic regions, and for the sequence from the negative sense molecule. Motifs for 40 human RBPs were found to be enriched in at least one of the tested genomic regions, while motifs for 23 human RBPs were found to be depleted in at least one of the tested regions (Supplementary Table 7).

We next examined whether any of the 6,936 putative binding sites for these 40 enriched RBPs were conserved across SARS-CoV-2 isolates. We found that 6,581 putative binding sites, representing 34 RBPs, were conserved across more than 95% of SARS-CoV-2 genome sequences in the GISAID database (≥ 26,213 out of 27,592 genomes). However, this is of limited significance as RBP binding sites in coding regions are likely to be conserved due to evolutionary pressure on protein sequences rather than RBP binding ability. We therefore repeated this analysis focusing only on putative RBP binding sites in the SARS-CoV-2 UTRs and intergenic regions. We found 124 putative RBP binding sites for 21 enriched RBPs in the UTRs and intergenic regions. Of these, 50 putative RBP binding sites for 17 RBPs were conserved in >95% of the available genome sequences; 6 in the 5’UTR, 5 in the 3’UTR, and 39 in intergenic regions (Supplementary Table 8).

Subsequently, we interrogated publicly available data to validate the putative SARS-CoV-2 / RBP interactions (Supplementary Table 9). According to GTEx data [25], 39 of the 40 enriched RBPs and all 23 of the depleted RBPs were expressed in human lung tissue. Further, 31 of 40 enriched RBPs and 22 of 23 depleted RBPs were co-expressed with the ACE2 and TMPRSS2 receptors in single-cell RNA-seq data from human lung cells (GSE122960; [25, 64]), indicating that they are present in cells that are susceptible to SARS-CoV-2 infection. We next checked whether any of these RBPs are known to interact with SARS-CoV-2 proteins and found that human poly-A binding proteins C1 and C4 (PABPC1 and PABPC4) bind to the viral N protein [24]. Thus, it is conceivable that these RBPs interact with both the SARS-CoV-2 RNA and proteins. Finally, we combined these results with our analysis of differential gene expression to identify SARS-CoV-2 interacting RBPs that also show expression changes upon infection. The results of this analysis are summarized for selected RBPs in Table 2.

**Table 2.** Selected conserved human RBPs predicted to interact with the SARS-CoV-2 genome along with experimental information.

| RBP | DE Analysis*1 |  |  | Experimental evidence in human datasets |  |  |  | RBP binding site prediction |  |
| --- | --- | --- | --- | --- | --- | --- | --- | --- | --- |
|  | A549 LogFC | Calu-3 LogFC | SARS-CoV-2 Specific DEG | GTEX Lung Tissue (TPM) | scRNA*2 | PPI Map*3 | Interaction with viral RNA*4 | Conserved*5 | Region |
| HNRNPA1 |  | -0.32 | ☐ | 331.336 | ✓ |  | ✓ | ✓ | 3'UTR |
| HNRNPA2B1 | -1.08 | -0.29 | ☐ | 539.829 | ✓ |  | ✓ | ✓ |  |
| PABPC1 | 0.72 | 0.44 | ✓ | 448.025 | ✓ | N | ✓ | ✓ |  |
| PABPC4 | 0.30 | -0.28 | ☐ | 103.082 | ✓ | N | ✓ | ✓ |  |
| PPIE |  | -0.27 | ☐ | 13.827 | ✓ |  | ☐ | ✓ |  |
| CELF5 | 0.56 |  | ☐ | 0.079 | ✓ |  | ☐ | ✓ | 5'UTR |
| FMR1 |  | 0.75 | ☐ | 21.435 | ✓ |  | ☐ | ✓ |  |
| RBM24 | 0.34 |  | ☐ | 1.412 | ☐ |  | ☐ | ✓ |  |
| EIF4B | 0.53 | 0.64 | ☐ | 170.303 | ✓ |  | ✓ | ✓ | Intergenic |
| ELAVL1 |  | -0.31 | ☐ | 27.440 | ✓ |  | ☐ | ✓ |  |
| PABPC1 | 0.72 | 0.44 | ✓ | 448.025 | ✓ | N | ✓ | ✓ |  |
| PPIE |  | -0.27 | ☐ | 13.827 | ✓ |  | ☐ | ✓ |  |
| TIA1 | 0.34 | 0.41 | ✓ | 46.934 | ✓ |  | ☐ | ✓ |  |
| TIAL1 | 0.25 |  | ☐ | 40.593 | ☐ |  | ☐ | ✓ |  |

\*1 LogFC reported only if padj < 0.05
\*2 scRNA expression in ACE+ and TMPRSS2+ lung cells: dataset GSE122960
\*3 PPI Map: Experimental map of protein-protein interactions between human and viral proteins (Gordon *et al.*, 2020)
\*4 Preprint: Experimental study revealing proteins interacting with SARS-CoV-2 RNA in a human liver cell line (Schmidt *et al.*, 2020)
\*5 Conserved in SARS-CoV-2 genomes

### Motif enrichment in SARS-CoV-2 differs from related coronaviruses

We repeated the above analysis to calculate the enrichment and depletion of RBP-binding motifs in the genomes of two related coronaviruses: the SARS-CoV virus (Supplementary Table 10) that caused the SARS outbreak in 2002-2003, and RaTG13 (Supplementary Table 11), a bat coronavirus with a genome that is 96% identical with that of SARS-CoV-2 [4, 88].

We found that the pattern of enrichment and depletion of RBP binding motifs in SARS-CoV-2 is different from that of the other two viruses. Specifically, the SARS-CoV-2 genome is uniquely enriched for binding sites of CELF5 in its 5’UTR, PPIE on its 3’UTR, and ELAVL1 in the viral negative-sense RNA molecule. These three proteins are involved in RNA metabolism and are important for RNA stability (ELAVL1, CELF5) and processing (PPIE). Despite the high sequence identity between the two genomes, the single binding site for CELF5 on the SARS-CoV-2 5’UTR is conserved in 97% of available SARS-CoV-2 genome sequences but absent in the 5’UTR of RaTG13.

### A subset of viral genome variants correlate with increased COVID-19 severity

To test whether any viral sequence variants were associated with a change in disease severity in human hosts, we analyzed 1,511 complete SARS-CoV-2 genomes that had associated clinical metadata. The FDR-corrected statistical results from this analysis revealed four nucleotide variations that were significantly associated with a change in viral pathogenesis. Three of these nucleotide changes resulted in non-synonymous variations at the amino acid level, while the last one was silent at the amino acid level. The first position was a *T*→*G* (L37F) substitution located in the Nsp6 coding region (p < 1.48E-5), the second position was a *C*→*T* (P323L) substitution located in the RNA-dependent RNA polymerase coding region (p < 2.01E-4), the third position was an *A*→*G* (D614G) substitution located in the spike coding region (p < 1.61E-4), and the fourth was a synonymous *C*→*T* substitution located in the Nsp3 coding region (p < 1.77E-4). As a further validation step, we performed the same analysis comparing viral sequence variants against potential confounders, such as the biological sex or age group of the patients. These comparisons validated that these four positions were only identified as significant in the results of the disease severity analysis.

## Discussion

Airway epithelial cells are the primary entry points for respiratory viruses and therefore constitute the first producers of inflammatory signals that, in addition to their antiviral activity, promote the initiation of the innate and adaptive immune responses. Here, we report the results of a complementary panel of analyses that enable a better understanding of host-pathogen interactions which contribute to SARS-CoV-2 replication and pathogenesis in the human respiratory system. Moreover, we propose already established along with novel human factors exclusively detected in SARS-CoV-2 infected cells by our analyses that might be relevant in the context of COVID-19 and which are worth being further investigated at an experimental level (Figure 4).

**Figure 4.**
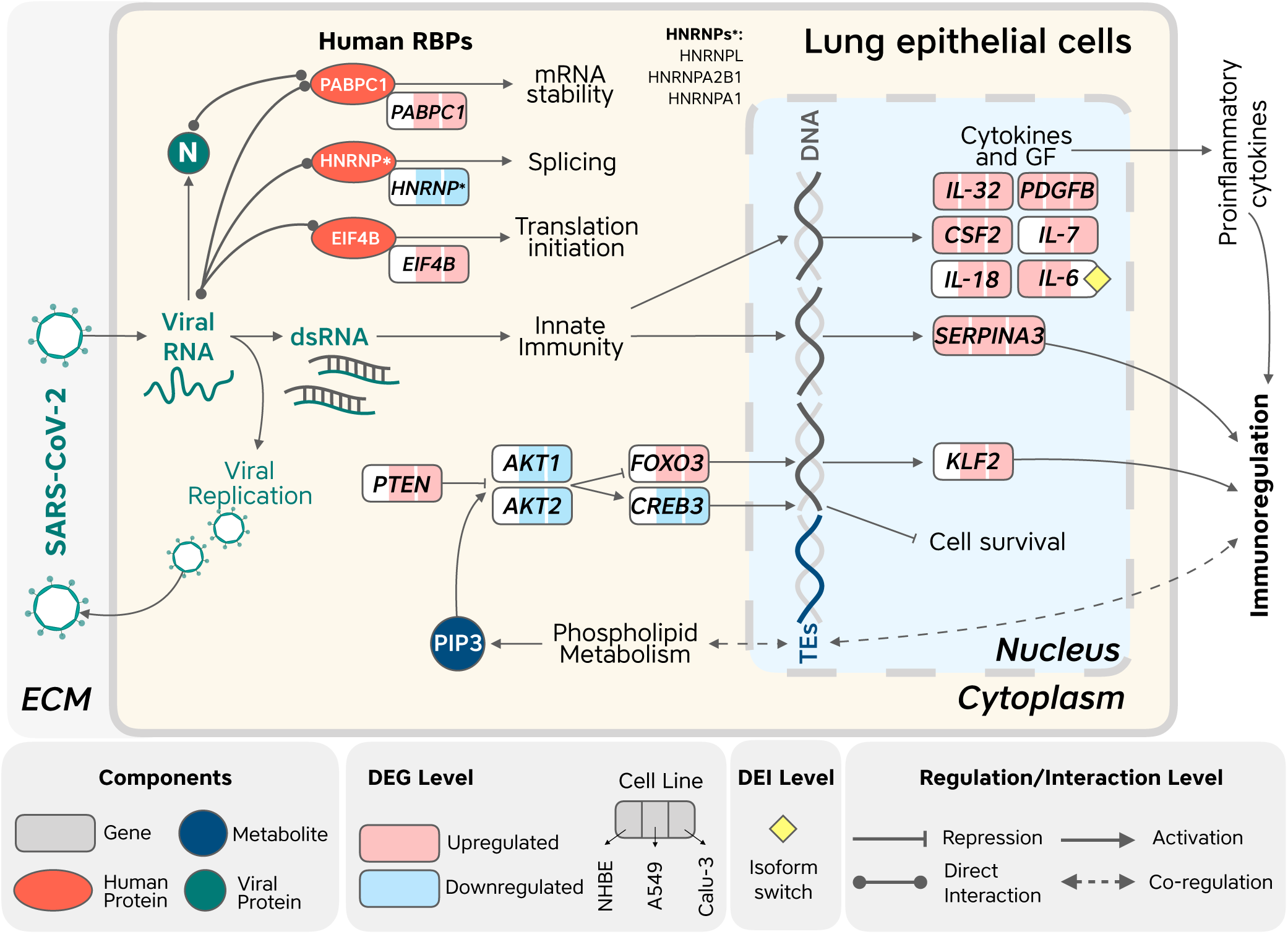
Overview of human factors specific to SARS-CoV-2 infection detected by our analyses. This includes human RBPs whose binding sites are enriched and conserved in the SARS-CoV-2 genome but not in the genomes of related viruses; and genes, isoforms and metabolites that are consistently altered in response to SARS-CoV-2 infection of lung epithelial cells but not in infection with the other tested viruses; ECM (extracellular matrix).

The *CSF2* gene, which encodes the Granulocyte-Macrophage Colony Stimulating Factor (GM-CSF), was among the most highly up-regulated genes in SARS-CoV-2 infected cells. GM-CSF induces survival and activation in mature myeloid cells such as macrophages and neutrophils. However, GM-CSF is considered more proinflammatory than other members of its family, such as G-CSF, and is associated with tissue hyper-inflammation [52]. In accordance with our results, high levels of GM-CSF were found in the blood of severe COVID-19 patients [84], and several clinical trials are planned using agents that either target GM-CSF or its receptor [53]. GM-CSF, together with other proinflammatory cytokines such as IL-6, TNF, IFNg, IL-7 and IL-18, is associated with the cytokine storm present in a hyperinflammatory disorder named hemophagocytic lymphohistiocytosis (HLH) which presents with organ failure [12]. Moreover, cytokines related to cytokine release syndrome (such as IL-1A/B, IL-6, IL-10, IL-18, and TNFA), showed increased positive association to the severity of the disease in the blood from COVID-19 patients [49]. Another proinflammatory cytokine specifically upregulated in SARS-CoV-2 infected cells was *IL-32*, which together with *CSF2*, promotes the release of TNF and IL-6 in a continuous positive loop and therefore contribute to this cytokine storm [90]. Interestingly, *IL-6, IL-7* and *IL-18* were found to be upregulated in two of the four data sets of SARS-CoV-2 infected cells. Moreover, not only upregulation, but also a shift in isoform usage of *IL-6* was detected in NHBE and A549 infected cells. A shift in 5’ UTR usage in the presence of SARS-CoV-2 may be attributed to indirect host cell signaling cascades that trigger changes in transcription and splicing activity, which could also explain the overall increase in *IL-6* expression.

*SERPINA3*, a gene coding for an essential enzyme in the regulation of leukocyte proteases, is also induced by cytokines [28]. This was the only gene consistently upregulated in all cell line samples infected with SARS-CoV-2 and absent from the other datasets. Even though this was previously proposed as a promising candidate for the inhibition of viral replication, to date no experiments were carried out to validate this hypothesis [13]. Another interesting candidate gene, which has not been implicated experimentally in respiratory viral infections and was upregulated in our analysis, was *VNN2*. Vanins are involved in proinflammatory and oxidative processes, and VNN2 plays a role in neutrophil migration by regulating b2 integrin [56]. In contrast, the downregulated genes included *SNX8*, which has been previously reported in RNA virus-triggered induction of antiviral genes [13, 26]; and *FKBP5*, a known regulator of NF-kB activity [27]. These results suggest that the SARS-CoV-2 virus tends to indirectly target specific genes involved in genome replication and host antiviral immune response without eliciting a global change in cellular transcript processing or protein production.

One of the first and most important antiviral responses is the production of type I Interferon (IFN). This protein induces the expression of hundreds of Interferon Stimulated Genes (ISGs), which in turn serve to limit virus spread and infection. Moreover, type I IFN can directly activate immune cells such as macrophages, dendritic cells and NK cells as well as induce the release of proinflammatory cytokines by other cell types [34]. Signaling pathway analysis showed that type I IFN response was greatly impacted in SARS-CoV-2 infected cells (A549 and Calu-3 cells at a MOI of 0.2 and 2 respectively). In the same direction, a higher expression of *PRDM1* (Blimp-1) that we observed in the SARS-CoV-2 infected cells, could also contribute to the critical regulation of IFN signaling cascades; interestingly, the TE family LTR13, which was also upregulated upon SARS-CoV-2 infection, is enriched in PRDM1 binding sites [78]. Therefore, it is possible that regulatory factors involved in IFN and immune response in the context of SARS-CoV-2 infection could also be attributed to TE transcriptional activation. In the same direction, we detected the upregulation of several TE families in SARS-CoV-2 infected cells that have been previously implicated in immune regulation. Moreover, 16 upregulated families were specific to SARS-CoV-2 infection in Calu-3 and A549 cell lines. The MER41B family, for instance, is known to contribute to interferon gamma inducible binding sites (bound by STAT1 and/or IRF1) [14]. Functional enrichment analysis of nearby genes were in accordance with these findings, since several immunity related terms were enriched along with “progressive pulmonary impairment”. In parallel, TEs seem to be co-regulated with phospholipid metabolism, which directly affects the Pi3K/AKT signaling pathway, central to the immune response and which were detected in our functional enrichment and metabolism flux analyses.

RBPs are another example of host regulatory factors involved either in the response of human cells to SARS-CoV-2 or in the manipulation of human machinery by the virus. We aimed at finding RBPs which potentially interact with SARS-CoV-2 genomes in a conserved and specific way. Five of the proteins predicted to be interacting with the viral genome by our pipeline (EIF4B, hnRNPA1, PABPC1, PABPC4, and YBX1) were experimentally shown to bind to SARS-CoV-2 RNA in an infected human liver cell line, based on a recent preprint [67].

Among the RBPs whose potential binding sites were enriched and conserved within the SARS-CoV-2 virus genomes is the EIF4B, suggesting that the SARS-CoV-2 virus protein translation could be EIF4B-dependent. We also detected the upregulation of *EIF4B* in A549 and Calu-3 cells, which might indicate that this protein is sequestered by the virus and therefore the cells need to increase its production. Moreover, this protein was predicted to interact specifically with the intergenic region upstream of the gene encoding the SARS-CoV-2 membrane (M) protein, one of four structural proteins from this virus.

Another conserved RBP, which was also upregulated in infected cells, is the Poly(A) Binding Protein Cytoplasmic 1 (PABPC1), which has well described cellular roles in mRNA stability and translation. PABPC1 has been previously implicated in multiple viral infections. The activity of PABPC1 is modulated to inhibit host protein transcript translation, promoting viral RNA access to the host cell translational machinery [72]. Importantly, the 3’ UTR region of SARS-CoV2 is also enriched in binding sites of the PABPC1 and the PPIE RBPs, the latter of which is known to be involved in multiple processes, including mRNA splicing [9, 33]. Interestingly, PABPC1 and PABPC4 interact with the SARS-CoV-2 N protein, which stabilizes the viral genome [24]. This raises the possibility that the viral genome, N protein, and human PABP proteins may participate in a joint protein-RNA complex that assists in viral genome stability, replication, and/or translation [1, 59, 62, 71, 72].

An interesting result was that the binding motifs for hnRNPA1, which has been shown to interact with other coronavirus genomes, were enriched specifically in the 3’UTR of SARS-CoV-2 even though they were depleted in the genome overall. The hnRNPA1 protein was described to interact more in particular with multiple sequence elements including the 3’UTR of the Murine Hepatitis Virus (MHV), and to participate in both transcription and replication of this virus [31, 44, 68]. This particular gene, along with *hnRNPA2B1*, were downregulated in Calu-3 cells and in contrast to the previous examples of upregulated genes, could denote a specific response of the human cells to control viral replication.

Cross referencing the results from our statistical analysis of ∼5% of the available genomes (∼1, 500 out of > 27,000 in GISAID) with clinical metadata revealed interesting new insights Indeed, the *D* → *G* mutation at amino acid position 614 in the Spike protein found in our analysis has recently been proposed to have increased viral infectivity [38]. In addition, this same mutation has also been associated with an increase in the case fatality rate [6], however, these hypotheses need further verification. The P323L mutation in the RNA-dependent RNA polymerase (RdRP) has been identified previously, although in that study it was associated with changes in geographical location of the viral strain [58]. Finally, the L37F mutation in the Nsp6 protein has been reported to be located outside of the transmembrane domain [11], being present at a high frequency [86], and proposed to negatively affect protein structure stability [8]. Our statistics may contain bias based on the number of genome sequences being collected earlier *versus* later in the pandemic, genomes lacking clinical outcome metadata, and in the case of the Spike D614G a potential increase of fitness associated with this mutation. However, the fact that more than one of our predictions has also been detected by different studies justifies future wet lab experiments to compare the effect of the other identified mutations.

## Conclusion

Overall, our analyses identified sets of statistically significant host genes, isoforms, regulatory elements, and other interactions that likely contribute to the cellular response during infection with SARS-CoV-2. Furthermore, we detected potential binding sites for human RBPs that are conserved across SARS-CoV-2 genomes, along with a subset of variants in the viral genome that correlate well with disease severity in SARS-CoV-2 infection. To our knowledge, this is the first work where a computational meta-analysis was performed to predict host factors that play a role in the specific pathogenesis of SARS-CoV-2, distinct from other respiratory viruses. Noteworthy, we used the only study available to date in which cells were infected with SARS-CoV-2 along with other viruses.

Other studies have focused on the sequencing of fluids (blood or bronchoalveolar lavage fluid), and may not reflect directly the cellular response to this virus. The availability of new high quality, well annotated datasets of patients and cells infected with SARS-CoV-2 will be valuable for the verification of our predictions. We envision that applying our publicly available workflow will yield important mechanistic insights in future analyses on emerging pathogens. Similarly, we expect that the results for SARS-CoV-2 will contribute to ongoing efforts in the selection of new drug targets and the development of more effective prophylactics and therapeutics to reduce virus infection and replication with minimal adverse effects on the human host.

## Materials and Methods

### Datasets

Two datasets were downloaded from the Gene Expression Omnibus (GEO) database, hosted at the National Center for Biotechnology Information (NCBI). The first dataset, GSE147507 [10], includes gene expression measurements from three cell lines derived from the human respiratory system (NHBE, A549, Calu-3) infected either with SARS-CoV-2, influenza A virus (IAV), respiratory syncytial virus (RSV), or human parainfluenza virus 3 (HPIV3). The second dataset, GSE150316, includes RNA-seq extracted from formalin fixed, paraffin embedded (FFPE) histological sections of lung biopsies from COVID-19 deceased patients and healthy individuals. This dataset encompasses a variable number of biopsies per subject, ranging from one to five. Given its limitations, we only utilized the second dataset for differential expression analysis.

The reference genome sequences of SARS-CoV-2 (NC_045512), RaTG13 (MN996532.1), and SARS-CoV (NC_004718.3) were downloaded from NCBI. Additionally, a list of known RBPs and their PWMs were downloaded from ATtRACT (https://attract.cnic.es/download). Finally, all SARS-CoV-2 complete genomes collected from humans and that had disease severity information were downloaded from GISAID on 19 May, 2020 [69].

### RNAseq data processing and differential expression analysis

Data was downloaded from SRA using sra-tools (v2.10.8; https://github.com/ncbi/sra-tools) and transformed to fastq with fastq-dump. FastQC (v0.11.9; https://github.com/s-andrews/FastQC) and MultiQC (v1.9) [20] were employed to assess the quality of the data used and the need to trim reads and/or remove adapters. Selected datasets were mapped to the human reference genome (GENCODE Release 19, GRCh37.p13) utilizing STAR (v2.7.3a) [17]. Alignment statistics were used to determine which datasets should be included in subsequent steps. Resulting SAM files were converted to BAM files employing samtools (v1.9) [43]. Next, read quantification was performed using StringTie (v2.1.1) [60] and the output data was postprocessed with an auxiliary Python script provided by the same developers to produce files ready for subsequent downstream analyses. For the second gene expression dataset, raw counts were downloaded from GEO. DESeq2 (v1.26.0) [47] was used in both cases to identify differentially expressed genes (DEGs). Finally, an exploratory data analysis was carried out based on the transformed values obtained after applying the variance stabilizing transformation [3] implemented in the vst() function of DESeq2 [48]. Hence, principal component analysis (PCA) was performed to evaluate the main sources of variation in the data and remove outliers.

### Gene ontology enrichment analysis

The DEGs produced by DESeq2 with an absolute Log2FC > 1 and FDR-adjusted p-value < 0.05 were used as input to a general GO enrichment analysis [5, 76]. Each term was verified with a hypergeometric test from the GOstats package (v2.54.0) [21] and the p-values were corrected for multiple-hypothesis testing employing the Bonferroni method [42]. GO terms with a significant adjusted p-value of less than 0.05 were reduced to representative non-redundant terms with the use of REVIGO [73].

### Host signaling pathway enrichment

The DEG lists produced by DESeq2 with an absolute Log2FC > 1 and FDR-adjusted p-value < 0.05 were used as input to the SPIA algorithm to identify significantly affected pathways from the R graphite library [65, 75]. Pathways with Bonferroni-adjusted p-values less than 0.05 were included in downstream analyses. The significant results for all comparisons from publicly available data from KEGG, Reactome, Panther, BioCarta, and NCI were then compiled to facilitate downstream comparison. Hypergeometric pathway enrichments were performed using the Database for Annotation, Visualization and Integrated Discovery (DAVID, v6.8) [30].

### Integration of transcriptomic analysis with the human metabolic network

To detect increased and decreased fluxes of metabolites we projected the transcriptomic data onto the human reconstructed metabolic network Recon (v2.04) [77]. First, we ran EBSeq [40] on the gene count matrix generated in the previous steps. Then, we used the output of EBSeq containing posterior probabilities of a gene being DE (PPDE) and the Log2FC as input to the Moomin method [63] using default parameters. Finally, we enumerated 250 topological solutions in order to construct a consensus solution for each of the datasets tested.

### Isoform Analysis

Using transcript quantification data from StringTie as input, we identified isoform switching events and their predicted functional consequences with the IsoformSwitchAnalyzeR R package (v1.11.3) [81]. In summary, we filtered for isoforms that experienced |≥30% |switch in usage for each gene and were corrected for false discovery rate (FDR) with a q-value < 0.05. Following filtering for significant isoforms, we externally predicted their coding capabilities, protein structure stability, peptide signaling, and shifts in protein domain usage using The Coding-Potential Assessment Tool (CPAT) [82], IUPred2 [18], SignalP [2] and Pfam tools respectively [19]. These external results were imported back into IsoformSwitchAnalyzeR and used to further identify alternative splicing events and functional consequences as well as visualize the overall and individual effects of isoform switching data. Specifically, to calculate differential analysis between samples, isoform expression and usage were measured by the isoform fraction (IF) value, which quantifies the individual isoform expression level relative to the parent gene’s expression level:

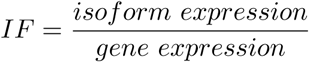

By proxy, the difference in isoform usage between samples (dIF) measures the effect size between conditions and is calculated as follows:

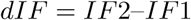

dIF was measured on a scale of 0 to 1, with 0 = no (0%) change in usage between conditions and 1 = complete (100%) change in usage. The sum of dIF values for all isoforms associated with one gene is equal to 1. Gene expression data was imported from the aforementioned DESeq2 results. The top 30 isoforms per dataset comparison were identified by ranking isoforms by gene switch q-value, i.e. the significance of the summation of all isoform switching events per gene between mock and infected conditions.

### Transposable Element Analysis

TE expression was quantified using the TEcount function from the TEtools software [41]. TEcount detects reads aligned against copies of each TE family annotated from the reference genome. DETEs in infected vs mock conditions were detected using DEseq2 (v1.26.0) [47] with a matrix of counts for genes and TE families as input. Functional enrichment of nearby genes (upstream 5kb and downstream 1kb of each TE copy within the human genome) was calculated with GREAT [51] using options “genome background” and “basal + extension”. We only selected occurrences statistically significant by region binomial test.

### Identification of putative binding sites for human RBPs on the SARS-CoV-2 genome

The list of RBPs downloaded from ATtRACT was filtered to human RBPs. The list was further filtered to retain PWMs obtained through competitive experiments and drop PWMs with very high entropy. This left 205 PWMs for 102 human RBPs. The SARS-CoV-2 reference genome sequence was scanned with the remaining PWMs using the TFBSTools R package (v1.20.0) [74]. A minimum score threshold of 90% was used to identify putative RBP binding sites.

### Enrichment analysis for putative RBP binding sites

The sequences of the SARS-CoV-2 genome, 5’UTR, 3’UTR, intergenic regions and negative sense molecule were each scrambled 1,000 times. Each of the 1,000 scrambled sequences was scanned for RBP binding sites as described above. The number of binding sites for each RBP was counted, and the mean and standard deviation of the number of sites was calculated for each RBP, per region, across all 1,000 simulations. A minimum FDR-adjusted p-value of 0.01 was taken as the cutoff for enrichment. This analysis was repeated with the reference genomes of SARS-CoV and RaTG13.

### Conservation analysis for putative RBP binding sites

The multiple sequence alignment of 27,592 SARS-CoV-2 genome sequences was downloaded from GISAID [69] on May 19th 2020. For each putative RBP binding site, we selected the corresponding columns of the multiple sequence alignment. We then counted the number of genomes in which the sequence was identical to that of the reference genome.

### Viral genotype-phenotype correlation

All complete SARS-CoV-2 genomes from GISAID, together with the GenBank reference sequence, were aligned with MAFFT (v7.464) within a high-performance computing environment using 1 thread and the –nomemsave parameter [55]. Sequences responsible for introducing excessive gaps in this initial alignment were then identified and removed, leaving 1,511 sequences that were then used to generate a new multiple sequence alignment. The disease severity metadata for these sequences was then normalized into four categories: severe, moderate, mild, and unknown. Next, the sequence data and associated metadata were used as input to the meta-CATS algorithm to identify aligned positions that contained significant differences in their base distribution between 2 or more disease severities [61]. The Benjamini-Hochberg multiple hypothesis correction was then applied to all positions [7]. The top 50 most significant positions were then evaluated against the annotated protein regions of the reference genome to determine their effect on amino acid sequence.

### Code availability

Code for these analyses is available at https://github.com/vaguiarpulido/covid19-research.

## Supporting Information

**Supplementary Figure 1.** Isoform Analysis.

**Supplementary File 1.** Zipped file containing Complete DEG tables.

**Supplementary File 2.** Zipped file containing GO for each dataset.

**Supplementary File 3.** TE family count/differential expression.

**Supplementary File 4.** GREAT Analysis (complete and per family).

**Supplementary Table 1.** Merged tables (Specific genes in SARS-CoV-2)

**Supplementary Table 2.** Supporting information for Figure 2, consisting of functional enrichment specific to SARS-CoV-2.

**Supplementary Table 3.** Pathway enrichment for each dataset (SPIA and DAVID merged into one file).

**Supplementary Table 4.** Metabolic fluxes predicted for each dataset using Moomin.

**Supplementary Table 5.** Isoform analysis.

**Supplementary Table 6.** Putative binding sites for human RBPs on the SARS-CoV-2 genome.

**Supplementary Table 7.** Enrichment of binding motifs for human RBPs on the SARS-CoV-2 genome.

**Supplementary Table 8.** Conservation of binding motifs for human RBPs across genome sequences of SARS-CoV-2 isolates.

**Supplementary Table 9.** Biological evidence associated with putative SARS-CoV-2 interacting human RBPs.

**Supplementary Table 10.** Enrichment of binding motifs for human RBPs on the SARS-CoV genome

**Supplementary Table 11.** Enrichment of binding motifs for human RBPs on the RaTG13 genome

## Acknowledgments

We would like to thank the Virtual BioHackathon on COVID-19 that took place during April 2020 (https://github.com/virtual-biohackathons/covid-19-bh20) for fostering an environment that triggered this collaboration and in particular the Gene Expression group for the fruitful discussions. This work was performed using the computing facilities of the CC/PRABI/LBBE in France and the France Génomique e-infrastructure (ANR-10-INBS-09-08), the HPC facilities of the University of Luxembourg [80]. We would also like to thank Slack for providing us with free access to the professional version of the platform.

We would also like to thank Slack for providing us with free access to the professional version of the platform.

